# Granular hydrogels as brittle yield stress fluids

**DOI:** 10.1101/2025.02.22.639638

**Authors:** G.B. Thompson, J. Lee, K.M. Kamani, N. Flores-Velasco, S.A. Rogers, B.A.C. Harley

## Abstract

While granular hydrogels are increasingly used in biomedical applications, methods to capture their rheological behavior generally consider shear-thinning and self-healing properties or produce ensemble metrics such as the dynamic moduli. Analytical approaches paired with common oscillatory shear tests can describe not only solid-like and fluid-like behavior of granular hydrogels but also transient characteristics inherent in yielding and unyielding processes. Combining oscillatory shear testing with consideration of Brittility (Bt) via the Kamani-Donley-Rogers (KDR) model, we show granular hydrogels behave as brittle yield stress fluids with complex transient rheology. We quantify steady and transient rheology as a function of microgel (composition; diameter) and granular (packing; droplet heterogeneity) assembly properties for mixtures of polyethylene glycol and gelatin microgels. The KDR model with Bt captures granular hydrogel behavior for a wide range of design parameters, reducing the complex transient rheology to a determination of model parameters. We describe the impact of composition on rheological behavior and model parameters in monolithic and mixed granular hydrogels. The model robustly captures self-healing behavior and reveals granular relaxation time depends on strain amplitude. This quantitative framework is an important step toward rational design of granular hydrogels for applications ranging from injection and *in situ* stabilization to 3D bioprinting.

## 1. Introduction

Hydrogels have enjoyed extensive use in biomedical applications in recent decades [1]. These hydrophilic polymer networks are excellent substrates for *in vitro* studies and *in vivo* applications due to their high capacity to store water, their biocompatibility, and the tunability of their biophysical properties such as diffusivity, to control release of cargo and biotransport, and viscoelasticity to mimic the mechanical behavior of a range of different tissues [2]. Formed from a liquid precursor, hydrogels are often stabilized with the introduction of physical or chemical crosslinks to prevent rapid dissolution. The use of synthetic polymers such as poly(ethylene) glycol (PEG) in cell-based applications typically necessitates the incorporation of bio-interactive crosslinkers that enable cellular adhesion and cell-mediated remodeling [3]. Hydrogels can also be formed from natural polymeric materials, including polysaccharides such as alginate and hyaluronic acid, or polypeptides such as silk fibroin, collagen, or gelatin [4]. These naturally-derived hydrogels present motifs that cells bind to or cleave, and have the potential to capture nonlinear mechanical behaviors typical of natural tissues [5]. While fabricated in many formats, hydrogels for biomedical applications have most commonly been deployed at scales (dimensions of order 10^-3^ m) significantly larger than that of cells (10^-5^ m).

By comparison, granular hydrogels are macrostructures formed by packing or jamming together hydrogel microparticles, or microgels [4]. The individual granular building blocks can be formed in a multitude of ways. Most common are mechanical fragmentation of a bulk hydrogel to produce irregularly shaped fragments [6, 7], or formation of spherical microgels via methods including electrospraying [8], microfluidic water-in-oil emulsification [9, 10], or batch mixing of an aqueous hydrogel phase in a continuous oil phase [11]. As a result, while granular hydrogels may be formed in similar volumes on a macrostructural basis with conventional hydrogels, their interstitial pore network enhances cellular infiltration, encourages network formation [12], and facilitates exchange of nutrients and waste between encapsulated cells and the surrounding environment [13]. Granular hydrogels are generally injectable [14] and further interlinking between microgels is not strictly necessary, even for *in vivo* applications [15]. Static covalent bonds [12], dynamic covalent bonds [16–18], or noncovalent stabilizing interactions [19, 20] may optionally be used to interlink individual microgels granting additional mechanical stability to granular scaffolds. Their multiscale properties enable a wide range of exciting application in biomedicine, including as templates to induce vascular integration and tissue repair *in vivo* [15, 21, 22] and as model systems to investigate multicellular interactions *in vitro* [4, 17, 23–25].

The mechanical behavior of viscoelastic materials such as hydrogels is often understood through the application of rheological techniques [26]. However, granular hydrogels can flow without suffering permanent degradation or fracture: large strains lead to the onset of flow and rearrangement within the macrostructure [27]. This reversible yielding process allows granular materials to behave as yield-stress fluids possessing the capability to transition from being soft solids at small stresses or strains to flowing like fluids at elevated stresses or strains [28, 29]. Improved understanding of the yielding behavior and nonlinear rheological performance is essential for a wide range of granular material applications, such as design of extrusion-based bioprinting of granular gels where large stresses or strains may be applied rapidly or sustained over an extended interval. Here, the ability to flow and re-solidify for structural integrity are necessary design criteria. Further, the yielding process can also significantly affect the viability and functionality of cells [30] embedded in the granular hydrogel.

Traditionally, the rheology of yield stress fluids has been conceptually broken into two distinct physical domains separated by the yield stress [31–35]. The unyielded behavior is commonly described by one equation with the physics of rigid, elastic, or viscoelastic solids [34], while the yielded behavior is described by another equation that represents the physics of a viscous fluid [36]. Within this Oldroyd-Prager formalism, as it has come to be known, yielding is thought of as an instantaneous transition that occurs as soon as the yield stress is exceeded [37, 38]. A proposal by Kamani, Donley, and Rogers, referred to here as the KDR model, presents yielding rheology as a gradual viscoelastic process that can be modeled at the continuum level [29]. The KDR model unifies the rheological physics of yield stress fluid by describing the yielded and unyielded behaviors within a single equation. Based on the concept of recovery rheology, the KDR model acknowledges the strain and strain rate as composite parameters made up of recoverable and unrecoverable components: *γ* = *γ_rec_* + *γ_unrec_* and *γ̇* = *γ̇_rec_* + *γ̇_unrec_* [39–42]. The model then accounts for yielding via a rate-dependent relaxation time. A recent extension of the model introduced the brittility factor, Bt, which affects the rate dependence of the relaxation time and allows for a description of a range of yielding behaviors from ductile to brittle responses [43].

In this work we show that the KDR model with brittility presents a holistic model of the rheology of granular hydrogels that provides predictive capabilities for *in situ* function and thereby reduces the complexity of the transient nonlinear rheology to a few design rules that dictate the model parameters. To date, experimental rheological data from granular hydrogels have been interpreted as shear-thinning, yielding, and self-healing behaviors, which have become seen as necessary properties for the use of granular hydrogels in a wide range of 3D printing applications. This effort provides a unified approach to consider the rheological behavior of homogenous and heterogenous granular assemblies, making it a highly flexible framework to describe the deformation and flow behavior of a broad range of granular hydrogels.

## 2. Materials and Methods

### 2.1 Synthesis and characterization of GelMAL

Gelatin maleimide (GelMAL) was synthesized as described previously [44]. Briefly, type A porcine gelatin, Bloom 300 (G1890, Sigma-Aldrich) was dissolved in a 4:5 (v/v) mixture of dimethyl sulfoxide (DMSO, MK494802, VWR) and 1 M 2-morpholinoethanesulfonic acid buffer (MES, Gold Biotechnology, St. Louis, MO) adjusted to pH 6.0. N-succinimidyl 3-maleimidopropionate (Tokyo Chemical Industry Co., Ltd) was dissolved in DMSO and added to the vial containing dissolved gelatin, the reaction mixture was adjusted to pH 4.5 with HCl, and the reaction was allowed to run for 24 hours at 40 °C. The resultant mixture was dialyzed in water at pH 3.25 at 40 °C for 5 days, frozen, lyophilized, and stored at -20 °C until use. The degree of functionalization of GelMAL was measured as described previously [44] by measuring relative to the phenylalanine peak using a ^1^H NMR spectrometer (Varian Unity Inova 400, 400 MHz).

### 2.2 Microgel formation through microfluidic emulsion

Microgels were formed as previously described using a flow-focusing microfluidic device with a 200 µm nozzle [10, 44]. An aqueous hydrogel precursor of either GelMAL or 4arm PEG Maleimide (PEG-4MAL, 20 kDa, JenKem Technology) dissolved in PBS, an oil mixture comprising light mineral oil (Sigma Aldrich) with 3% SPAN80 (Sigma Aldrich), and an emulsion of 30 mg/mL DTT added to the aforementioned oil mixture at a ratio of 1:12.6 DTT solution:oil. Solutions were pumped into a microfluidic device using syringe pumps (Pump 11 Elite, Harvard Apparatus). For 140-µm microgels, the flowrates were 10 µL/min, 1 µL/min, and 80 µL/min for the hydrogel precursor, oil, and crosslinker emulsion, respectively. Microgels were collected in PBS and placed on a shaker at 4 °C for 20 minutes. Then, acellular microgels were washed in PBS + 0.1% Tween-20 (Fisher Scientific) to remove residual oil, followed by three washes in PBS. Washes were performed at 600 rcf x 5 minutes.

### 2.3 Microgel formation through batch emulsion

The batch emulsion method required 3 different mixtures: GelMAL precursor, light mineral oil + 3 vol% SPAN80, and DTT emulsion in ratio of 1:30:6 (v/v). The GelMAL to 3 vol% SPAN80 light mineral oil ratio was inspired by batch emulsion work performed by others [11] and the GelMAL to DTT emulsion ratio of 1:6 was chosen to match the flow ratio of the components when using microfluidics. The GelMAL solution was created by weighing out GelMAL and dissolving in PBS on a hot plate set to 43 °C with vigorous stirring to create a 4 wt% solution. The DTT emulsion was created by weighing out the DTT; dissolving it in PBS to create a 30 mg/mL solution; adding it, in a ratio of 1:12.6 (v/v), to a separate solution of light mineral oil + 3 vol% SPAN80; and then emulsifying by sonication. The batch emulsion apparatus was a separate hot plate with an aluminum heating block, a thermometer, and a 20mL scintillation vial. The 3 vol% SPAN80 light mineral oil was transferred to the scintillation vial and heated with stirring at the desired agitation rate until the oil reached 38 °C. GelMAL solution was then pipetted dropwise into the vial and was allowed to be sheared into microdroplets for 2 minutes. After two minutes, the DTT emulsion was pipetted dropwise into the vial and then allowed to mix for two minutes. Afterwards, the contents of the vial were transferred to a centrifuge tube and washed in PBS + 0.1% Tween-20 to remove residual oil. Finally, microgels were washed three times in PBS at 300 rcf x 5 min.

### 2.4 Quantification of microgel diameter

Brightfield images of microgels were taken with a DMi8 Yokogawa spinning disk confocal microscope fitted with a Hamamatsu EM-CCD digital camera (Leica Microsystems, Inc., Deerfield, IL). Diameters were measured using ImageJ. A minimum of 100 microgels were measured to describe the output of the flow condition.

### 2.5 Granular hydrogel formation

Granular hydrogels were formed through a two-step process. First, washed microgels were centrifuged at 300 rcf x 5 min to achieve a moderate packing density and the supernatant was aspirated. Then, a filter (GVWP04700, Millipore Sigma) was formed into a cone and placed in a 15-mL conical. Microgels were transferred to the filter cone via pipette and the sample was centrifuged for 300 rcf x 30 s to achieve a high packing density (Fig 1A) [47].

**Figure 1.**
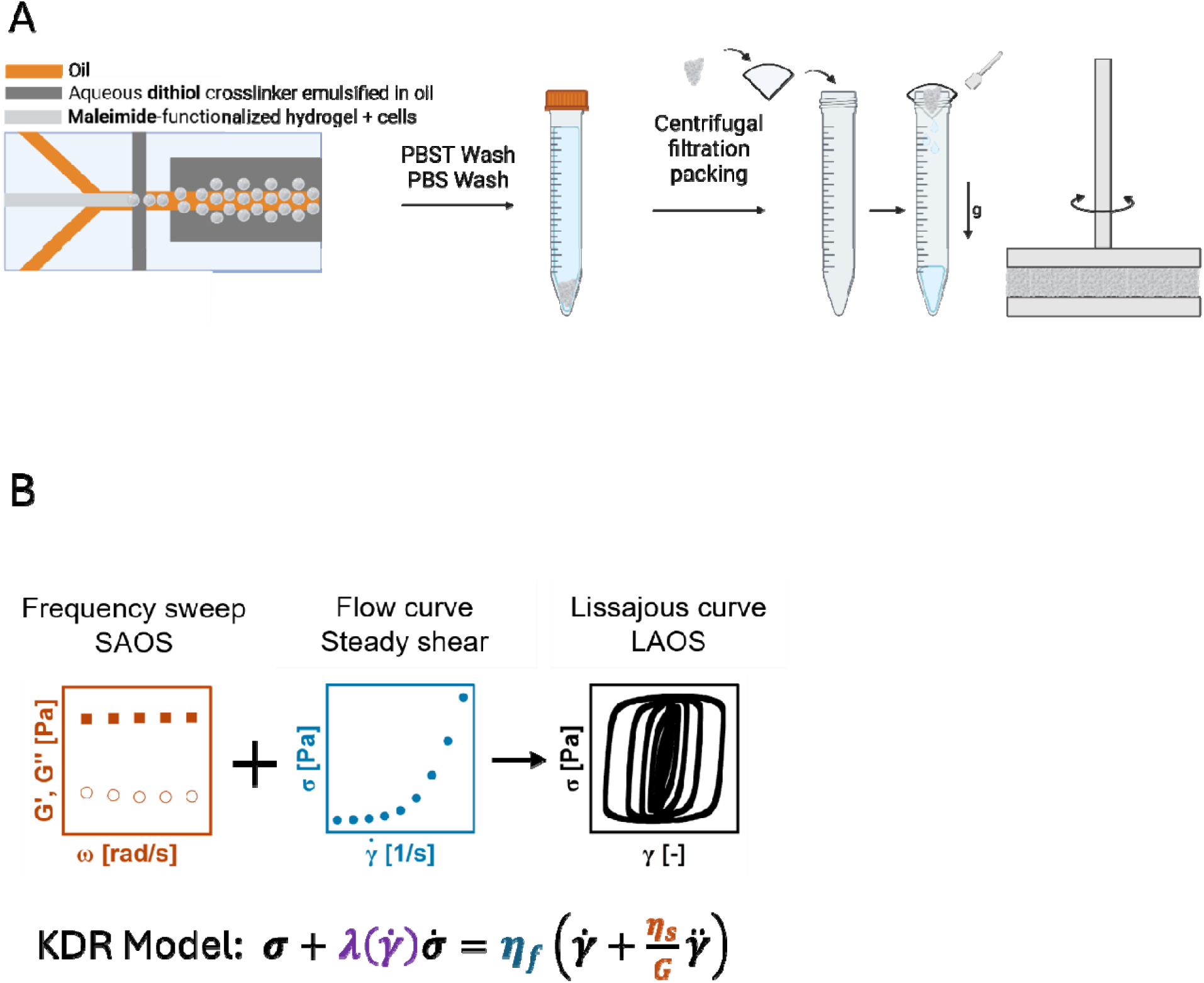
**A** Overview of microgel fabrication and granular hydrogel formation process. Microgels are formed using a flow-focusing microfluidic device. Microgels are then washed with PBST and PBS to remove oil and residual surfactant, respectively. Microgels are packed into a granular gel by transferring to a filter placed at the top of a conical and centrifuged. Finally, microgels are transferred to the rheometer for characterization. **B** Description of modeling framework. In the KDR model, the elastic modulus *G* and structural viscosity *η_s_* are inferred from a frequency sweep. The consistency index *k*, flow index *n*, and yield stress *σ_y_* are taken from the Herschel-Bulkley fit to the characteristic flow curve. In this study, brittility *Bt* is a fitted parameter that represents the ratio of the local modulus to the bulk modulus of the macrostructure. These parameters are fed to the KDR model, which can predict the stress response to an induced strain.

### 2.6 Rheological testing

All granular hydrogels were tested within four hours of microgel formation and packing. Tests were performed on an Anton Paar Modular Compact Rheometer (MCR) 302 using 20mm parallel plate geometry. Granular hydrogels were scooped from the filters and onto the rheometer using a spatula.

All tests are carried out using oscillatory shearing. Linear viscoelastic properties were determined by sweeping over frequencies from 1 to 10 rad/s at a small amplitude of 1%. These were repeated between each nonlinear test to ensure reproducibility. Nonlinear responses were determined from amplitude sweeps carried out at 1 rad/s, with amplitudes ranging from 0.1% up to 1000%. Self-healing tests were carried out across a range of large and small amplitudes, starting at small amplitudes, stepping up to large amplitudes, and then reducing the amplitude back to the original small value [48].

### 2.7 Kamani-Donley-Rogers model and parameter determination

The KDR model describes the rheology of yield stress fluids above and below the yield stress within the same continuous framework [29]. While the full version of the model is tensorial, the 1-D version, which describes the shear component of the stress-strain relation is

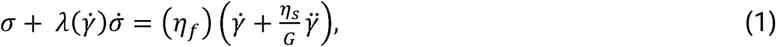

where *σ* and *σ̇* are the shear stress and its derivative, *γ̇* and *γ̈* are the strain rate and its derivate, *λ*(*γ̇*) is the rate-dependent relaxation time, *η_f_* is the flow viscosity, and *η_s_* and *G* are the structural viscosity and elastic modulus. The model is constructed on the basis that strain is a composite parameter that can be experimentally divided into recoverable and unrecoverable components. Similarly, the KDR model can be divided into recoverable and unrecoverable behaviors to determine the parameters.

When subjected to small stresses or strains, yield stress fluids behave like viscoelastic solids whose rheological response is predominantly recoverable [29]. At large stresses or strains, they behave like fluids whose rheology is predominantly unrecoverable. The amplitude sweep is an experimental protocol that encompasses these two behaviors and is therefore highly useful in determining the model parameters [49].

The KDR model has 6 parameters that need to be determined. The elastic modulus, *G*, and structural viscosity, *η_s_*, can be determined from the linear viscoelastic response elicited at small amplitudes. In the linear viscoelastic regime, the storage modulus, *G*′(*ω*), and loss modulus, *G*″(*ω*), are weakly dependent on the frequency. The recoverable behavior of the model can therefore be specified by *G* = *G*′(*ω*) and, *η_s_* = *G*″(*ω*)/*ω*, from either a frequency sweep in the linear regime or the small amplitude response of the amplitude sweep. For consistency, we determine *G* and *η_s_* at *ω* = 1 rad/s.

The KDR model describes the unrecoverable, or flow behavior, with the form of a Herschel-Bulkley fluid with an effective shear rate that depends on the brittility,

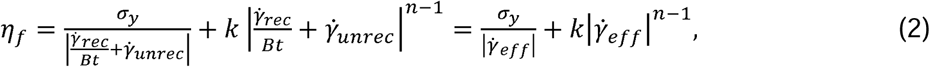

where *σ_y_* is the yield stress, *k* is the consistency index, *n* is an exponent that describes the high shear rate flow behavior, and *Bt* is the Brittility factor [43]. As discussed above, *γ̇_rec_* and *γ̇_unrec_* are the rates at which strain is acquired recoverably and unrecoverably. Equation 2 succinctly expresses the major contribution of the KDR model: the unrecoverable flow behavior is dependent on the recoverable elastic response. The KDR model therefore describes yielding as a nonlocal process where elastic deformation in part of the material assists yielding and plastic behavior somewhere else. The formulation of the model is agnostic to spatial dependence, meaning that the description is a mean-field only. The allowance for plastic behavior to be assisted by elastic behavior stands in stark contrast to other descriptions of yield stress fluids, which describe the flow behavior in terms of unrecoverable strains rates only.

The yielding behavior is described by the KDR model via the rate dependent relaxation time, λ(*γ̇*), which combines the recoverable and unrecoverable components,

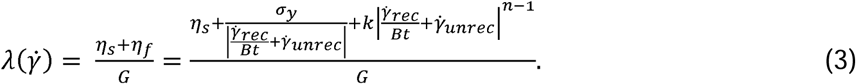

While the elastic modulus and structural viscosity can be determined from the small amplitude response, *σ_y_*, *k*, *n*, and *Bt*, can be determined by fitting the transient flow behavior at the largest amplitudes. The dynamic moduli, *G*′ and *G*″, which represent the average amount of energy stored and dissipated, are given by,

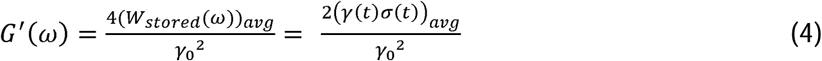

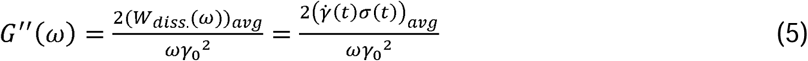

To construct a full sweep, the steady state response is calculated for a range of applied amplitudes. All 6 model parameters can therefore be obtained using experimental data from a combination of frequency and amplitude sweeps, or from an amplitude sweep alone, including the transient amplitude sweep data (Fig 1B). Once the model parameters have been obtained, predictions of the material behavior under arbitrary deformations or loading protocols can be made.

### 2.8 Statistics

Microgel size statistics were reported as the mean, standard deviation (SD), and coefficient of variation (% CV, standard deviation divided by the mean). Stiffness is reported as the mean ± standard error of the mean. Normality was evaluated using the Shapiro-Wilkes test [50, 51] at a 0.05 significance level. Conditions were compared using the Kruskal-Wallis test [52] and the Wilcoxon Rank Sum test [53, 54]. All plots were created using OriginPro 2024.

## 3. Results

### 3.1 Granular gel porosity is sensitive to microgel properties and granular gel preparation

Compared to conventional bulk hydrogels, granular hydrogels possess an expanded space of tunable design parameters such as microgel structural and mechanical features as well as the packing and diversity of microgel populations used to form the granular gel. We first formed microgels from 4wt% GelMAL and 5wt% PEG-4MAL to create heterogeneous granular mixtures from two mechanically distinct materials. We formed 4wt% GelMAL and 5wt% PEG-4MAL microgels using a flow focusing microfluidic device (MD) with a flowrate ratio of 8:1 (continuous phase:hydrogel phase). These conditions yielded microgels of similar, but statistically different size (Fig 2A-B, Table 1). Then, taking 5 wt% PEG-4MAL microgels as a reference material, we formed additional PEG-4MAL microgel variants in which fabrication parameters were altered to define the effect of these factors on granular hydrogel rheology and yielding behavior (Fig 2A-B, Table 1). To investigate the effect of microgel size on granular properties, we decreased the flowrate ratio from 8:1 to 4:1, increasing the microgel diameter from ∼140 µm to ∼200 µm. Lowering the wt% of PEG-4MAL from 5% to 4% also slightly reduced microgel diameter. To achieve a microgel population with an equivalent average diameter but higher variability in microgel size, we used batch emulsification. Here, after screening a range of stirring rates for the batch emulsification process (Table S1), we identified a stirring rate (300 RPM) that generated microgels of equivalent average diameter to the microfluidic device while increasing the CV in microgel diameter from 4.1% to 29.7% (Table 1).

**Figure 2.**
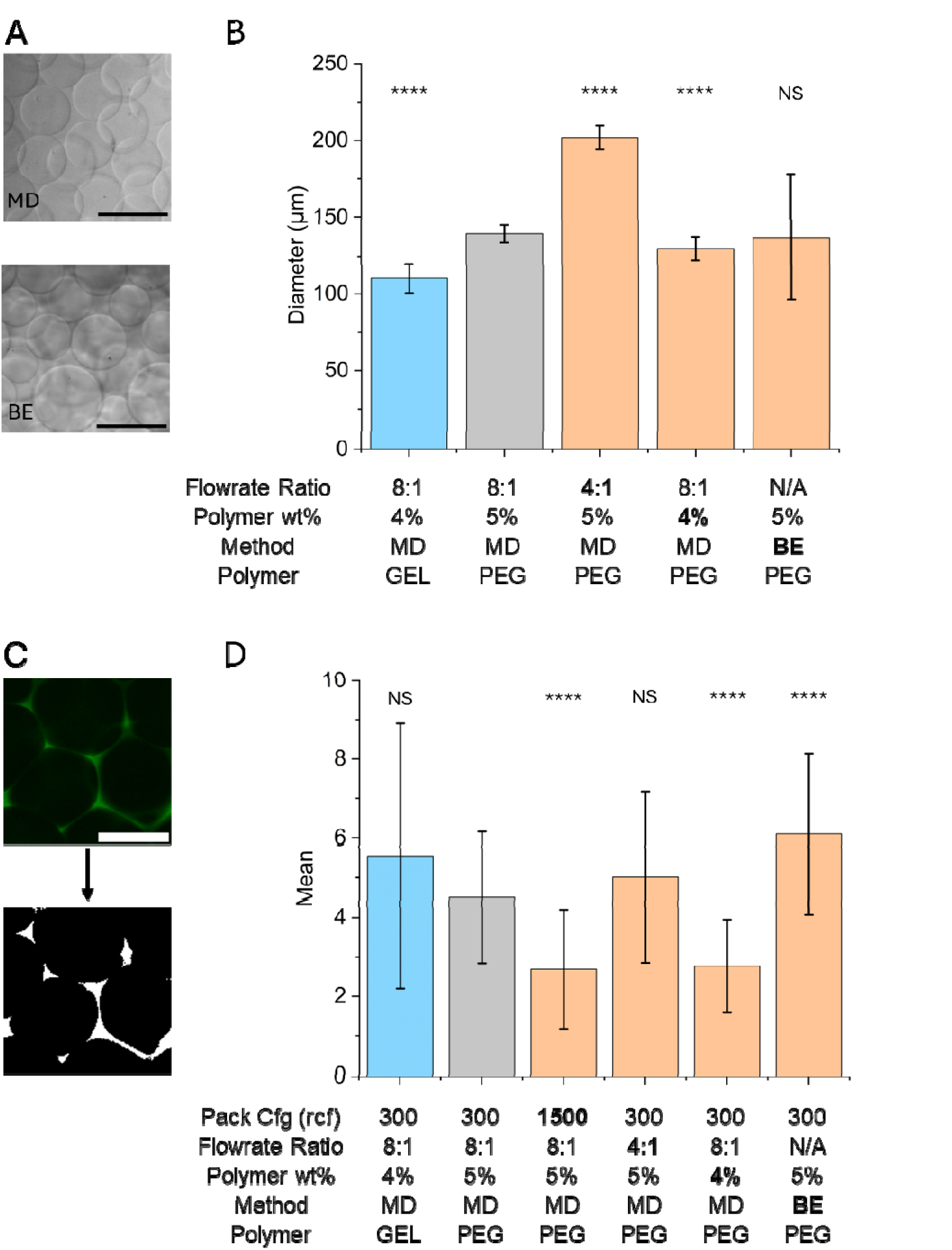
**A** Microgels formed using microfluidics (top) and batch emulsion (bottom) imaged under brightfield. **B** Microgel diameter for the range of key conditions tested in this study. Data are reported as mean ± standard deviation with n > 100 microgels per condition. Statistical significance is reported relative to the reference condition (Small / 5% / MFD / PEG). **C** Fluorescent imaging and thresholding for granular gel. **D** Granular hydrogel porosity for the range of key conditions tested in this study. Data are reported as mean ± standard deviation with n = 60 z-slices (20 from each of 3 granular gels) per condition. Statistical significance is reported relative to the reference condition (Low / Small / 5% / MFD / PEG).

**Table 1.** Microgel diameter statistics for key microgel populations used in this study. MFD = flow focusing microfluidic device, BE = batch emulsion.

| **Microgel Size** | **Small** | **Large** | **Small** | **Small** | **Small** |
| --- | --- | --- | --- | --- | --- |
| **Polymer wt%** | 5% | 5% | 4% | 5% | 4% |
| **Method** | MFD | MFD | MFD | BE | MFD |
| **Polymer** | PEG-4MAL | PEG-4MAL | PEG-4MAL | PEG-4MAL | GeIMAL |
| Mean ($\mu\text{m}$) | 139.7 | 201.9 | 130.0 | 137.2 | 110.3 |
| SD ($\mu\text{m}$) | 5.7 | 7.7 | 7.7 | 40.8 | 9.7 |
| % CV | 4.1% | 3.8% | 5.9% | 29.7% | 8.8% |

We subsequently defined the effect of centrifugal force used for packing the granular gels on the porosity of the resulting granular hydrogels for the group of microgel variants (Fig 2C-D, Table 2). Despite shifts in microgel size, we observed that granular gels formed from 4wt% GelMAL and 5wt% PEG-4MAL displayed similar porosities. Notably, increasing microgel size did not significantly alter porosity of the resultant granular gel in 5wt% PEG-4MAL variants. However, increasing the centrifugal force in the final preparation step from 300g to 1500g reduced porosity from 4.5% to 2.7%. Decreasing the wt% of PEG-4MAL similarly reduced porosity to 2.8%. Finally, granular hydrogels formed form more irregular microgels created via batch emulsion significantly increased porosity (6.1%). Together, these results indicate that tuning microgel design parameters, microgel fabrication schemes, and granular hydrogel packing methods can significantly alter the available open network spaces (µm-scale properties) of granular assemblies.

**Table 2.** Granular hydrogel porosity statistics for key granular hydrogels explored in this study.

|  |  |  |  |  |  |  |
| --- | --- | --- | --- | --- | --- | --- |
| **Packing** | **Low** | **High** | **Low** | **Low** | **Low** | **Low** |
| **Microgel Size** | **Small** | **Small** | **Large** | **Small** | **Small** | **Small** |
| **Polymer wt%** | **5%** | **5%** | **5%** | **4%** | **5%** | **4%** |
| **Method** | **MFD** | **MFD** | **MFD** | **MFD** | **BE** | **MFD** |
| **Polymer** | **PEG-4MAL** | **PEG-4MAL** | **PEG-4MAL** | **PEG-4MAL** | **PEG-4MAL** | **GeIMAL** |
| Mean (%) | 4.50 | 2.69 | 5.01 | 2.77 | 6.10 | 5.55 |
| SD (%) | 1.66 | 1.51 | 2.16 | 1.18 | 2.03 | 3.35 |

### 3.2 Rheology of granular gel composites formed from discrete microgel populations

Having characterized geometric properties of a library of microgels and granular hydrogels, we next evaluated the rheological performance of granular hydrogels formed purely from 4wt% GelMAL microgels, purely from 5wt% PEG-4MAL microgels, or mixtures (2:1 and 1:2, GelMAL:PEG-4MAL) of these two microgel populations. Amplitude sweeps, (Fig 3A, top row) reveal changes in both linear and nonlinear viscoelastic behavior as the fraction of PEG-4MAL is increased in the granular composite. Lissajous curves (Fig 3B, top row) depict the transient shear rheology, with the shear stress plotted parametrically against the shear strain over one period. The KDR model with Bt is used to quantitatively assess rheological behavior in the granular hydrogels and to reduce the complexity of the transient nonlinear rheology to a dependence of the model constitutive properties on granular composition. The model accurately captures the linear and nonlinear viscoelastic behaviors of the granular hydrogels composed of two discrete microgel populations (Fig 3A & 3B; bottom rows). Notably, as the fraction of PEG-4MAL within the granular gel increases, so too do the magnitude of the recoverable properties characterized by the elastic modulus and structural viscosity (Fig 3C, Table S2). The yielding of granular gels composed primarily or entirely of PEG-4MAL is much more brittle. This phenomenon can be qualitatively assessed by the more rapid increase in the loss modulus at intermediate strain amplitudes (Fig 3A (e,g)), as well as the buildup and sudden decrease of stress as the material yields, as shown in the top-left and bottom-right of each Lissajous curve at large strain amplitudes (Fig 3B (e,g)). These observations are associated with an increase in Bt from 2 to 8 as the fraction of PEG-4MAL in the granular gel increases from 0% to 100% (Fig 3C). The model yield stress values vary with the PEG-4MAL fraction nonmonotonically.

**Figure 3.**
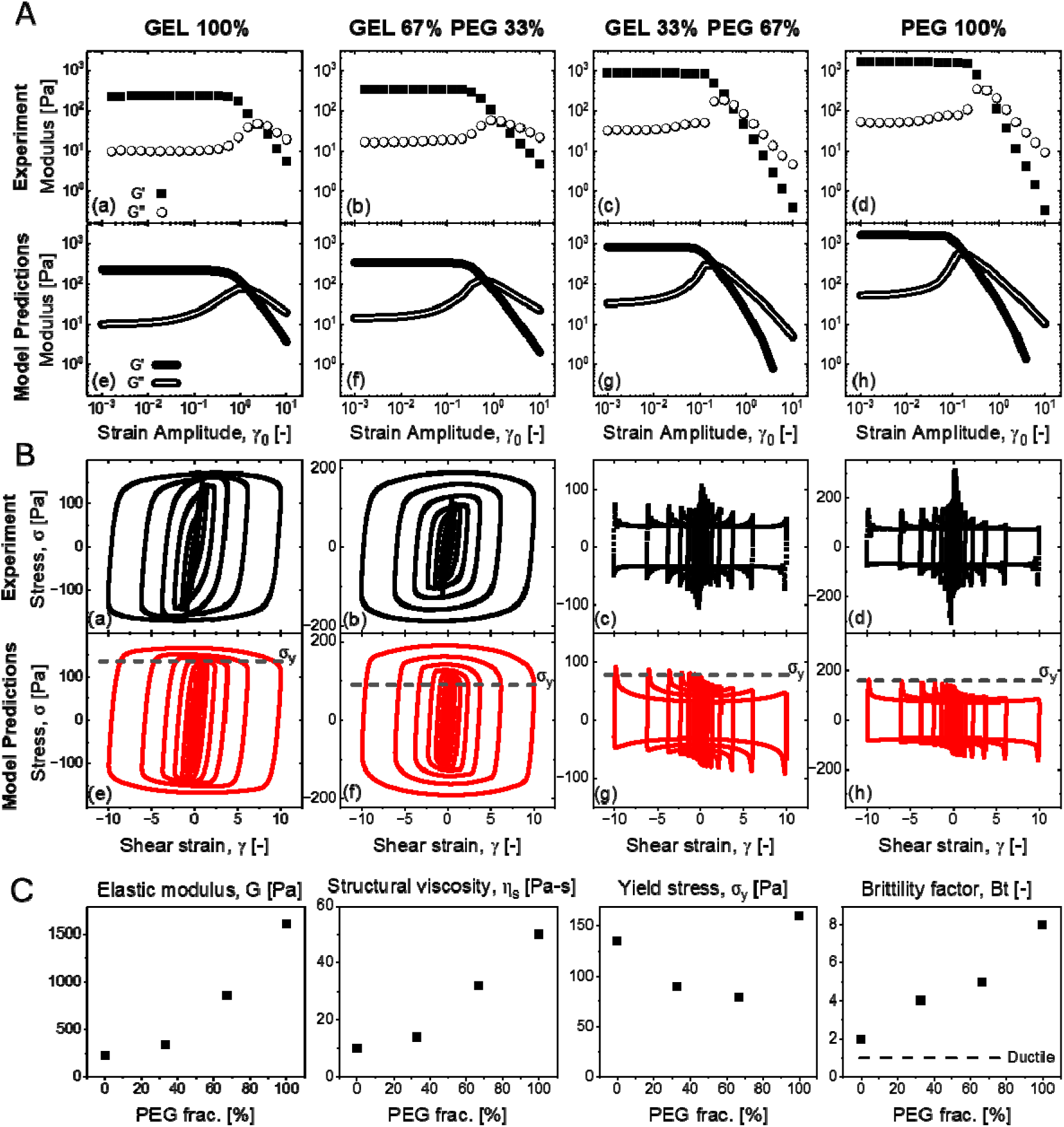
**A** Experimental (a,b,c,d) and KDR model-predicted (e,f,g,h) amplitude sweeps for heterogeneous granular hydrogels comprising a range of fractions of 4wt% GelMAL and 5wt% PEG-4MAL microgels. Storage moduli (G’) are represented with darkened points or lines, and loss moduli (G’’) are represented by open points or lines. **B** Experimental (a,b,c,d) and KDR model-predicted (e,f,g,h) Lissajous curves from the amplitude sweep at 1 rad/s, showing the full range of behavior under oscillatory shear for heterogeneous granular hydrogels comprising a range of fractions of 4wt% GelMAL and 5wt% PEG-4MAL microgels. Dotted lines indicate the fitted yield stress, *σ_y_*, for each composite. **C** Scatterplots depict the fitted elastic modulus *G,* structural viscosity *η_s_*, yield stress *σ_y_,* and brittility *Bt* for each granular hydrogel, as a function of the fraction of 5wt% PEG-4MAL microgels. The dotted line on the brittility plot indicates a perfectly ductile material with a corresponding *Br* of 1.

A change in the yielding behavior is clearly observed between the 33% and 67% PEG-4MAL conditions. The shape of the Lissajous curves at large strain amplitudes change from being smoothly rounded to having sharp yielding transitions (Fig 3B (c,e)). When the stiffer PEG-4MAL dominates the microgel population, there is a sudden decrease of stress when the material yields and plastic flow dominates (Fig 3B (e,g)). At the largest strain amplitudes, the steady stress values during the plastic flow, indicated by the horizontal flat regions in the Lissajous curves, exceeds 150Pa for the soft GelMAL-dominated granular hydrogels, while the steady stress values of stiffer PEG-4MAL-dominated granular hydrogels are less than 100 Pa. These observations indicate that the linear and nonlinear viscoelastic properties of heterogeneous granular hydrogels can be tuned by varying the fraction of each microgel population. Further, our observations suggest the possibility that the discrete mixtures of heterogenous microgels can be revealed from linear and nonlinear viscoelastic properties recovered from the KDR model.

### 3.3 Impact of microgel characteristics and granular properties upon granular gel rheology

The impact of tunable fabrication parameters on granular mechanics was explored using the 5wt% PEG-4MAL as a reference condition (Fig 4A-B). The tunable parameters include the microgel diameter, weight percentage (wt%), packing density and processing method. Increasing the microgel diameter from ∼140 µm to ∼200 µm had little effect on elastic modulus but decreased the structural viscosity. While there was an increase in the yield stress, the brittillity remained the same. Similarly, reducing the porosity from 4.5% to 2.5% (increasing packing density) had negligible effects on the elastic modulus but led to a similar decrease in the structural viscosity. While the brittillity did not change with decreased porosity, there was a significant increase in the yield stress. The increased %CV in particle diameter achieved via batch processing led to a modest decrease in the elastic modulus as well as a similar decrease in structural viscosity. The yield stress and the brittility factor for these more heterogenous granular assemblies also decreased as the overshoot in G’’ became less abrupt and reflected less brittle yielding behavior. Interestingly, decreasing polymer weight percent of the hydrogel microparticles from 5% to 4% induced marked changes of all the properties. Both recoverable components, the elastic modulus and structural viscosity, and the yield stress significantly decreased, whereas the brittility factor increased substantially. This indicates that the material became much softer, but the yielding behavior became more brittle, meaning the nonlinear behavior is more rapid. The effects of each granular hydrogel design parameter upon model parameters are summarized in Table S3. A comparison may also be made here between PEG-4MAL and GelMAL microparticles held at a constant polymer wt%. In this case, 4wt% PEG-4MAL possessed a greater elastic modulus, greater structural viscosity, and greater brittility (20 vs 2) than 4wt% GelMAL. Together, these parameters reveal that the PEG-4MAL granular assembly is a stiffer and more brittle material than GelMAL at the same polymer fraction.

**Figure 4.**
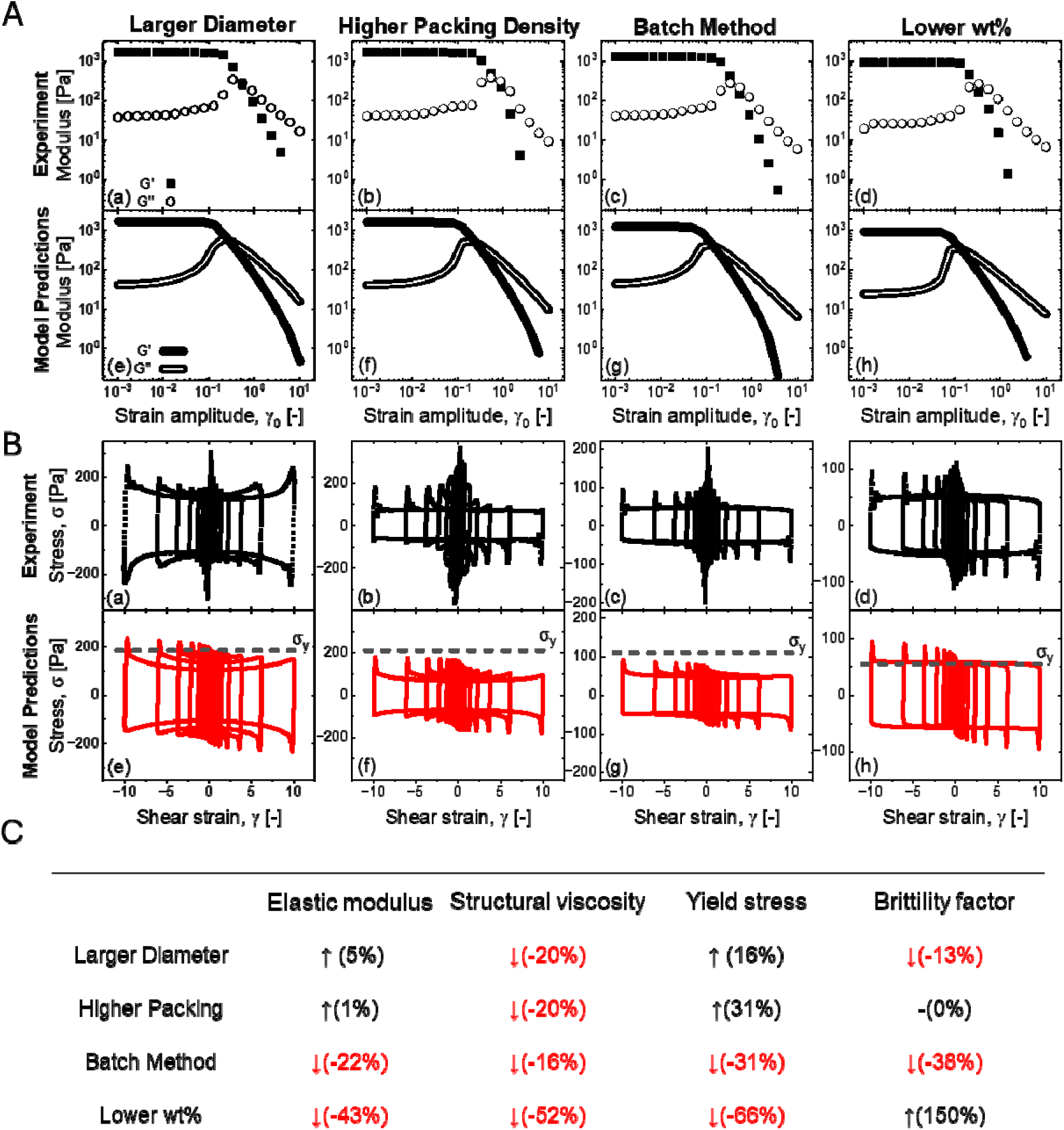
**A** Experimental (a,b,c,d) and KDR model-predicted (e,f,g,h) amplitude sweeps for PEG-4MAL granular hydrogels made with altered granular hydrogel parameters of microgel diameter, granular hydrogel packing density, microgel formation method, and microgel polymer fraction. Storage moduli (G’) are represented with darkened points or lines, and loss moduli (G’’) are represented by open points or lines. **B** Experimental (a,b,c,d) and KDR model-predicted (e,f,g,h) Lissajous curves from the amplitude sweep at 1 rad/s, showing the full range of behavior under oscillatory shear PEG-4MAL granular hydrogels made with altered granular hydrogel parameters of microgel diameter, granular hydrogel packing density, microgel formation method, and microgel polymer fraction. Dotted lines indicate the fitted yield stress, *σ_y_*, for each composite. **C** Trends observed in fitted parameters used to describe granular hydrogel behavior in monolithic PEG-4MAL granular hydrogel variations.

### 3.4 Self-healing

Extrusion based 3D printing techniques often involve highly varied force dynamics. When loaded in the barrel prior to printing, forces on the material are low. During the extrusion process, forces are high, but they drop again once the material exits the nozzle. Depending on the shape of the printed object, and the location within that shape, material forces after that point may then be close to zero or remain significant but static. The ability of the material to regain the mechanical properties exhibited before being made to flow is referred to as ‘self-healing’. This self-healing behavior is quantified by a rheological test in which the oscillatory shear applied alternates between small and large amplitudes. The precise meaning of low and high stress is determined through the amplitude sweep. Low stress is typically considered to be any amplitude in the linear regime, before yielding has taken place. High stress can be any amplitude that elicits a nonlinear response, but it is common to select an amplitude beyond which the dynamic moduli have crossed to ensure yielding takes place. In our case, we selected strain amplitudes of 0.01 and 0.1 strain units as representative ‘small’ amplitudes, so that we can also assess the effects of the rate-dependent relaxation time on the self-healing behavior. The experimental data obtained from self-healing tests exhibits a complete recovery of G’ and G’’ after the high stress application. Predictions of the KDR model with Bt also capture the transient changes of the dynamic moduli and its complete recovery (Fig 5, Fig S1).

**Figure 5.**
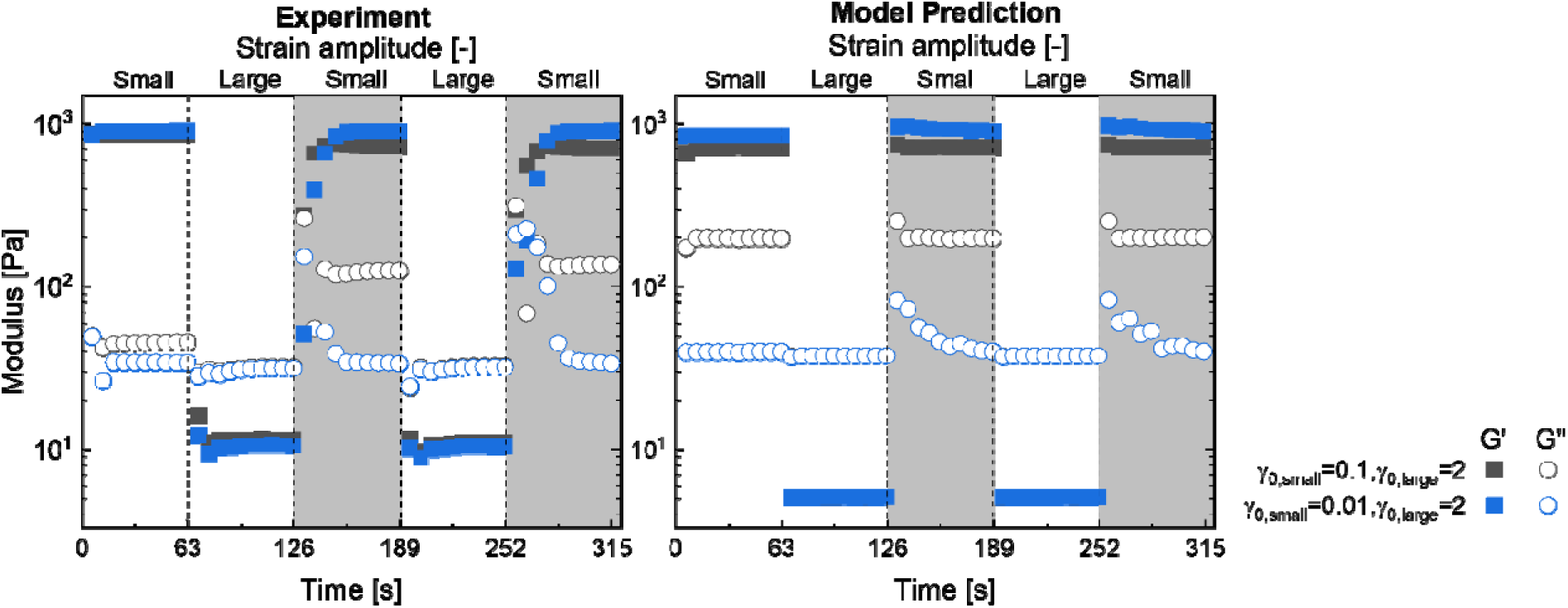
Experimental (left) and KDR model-predicted (right) self-healing data for granular hydrogel mixtures 1:2 GelMAL:PEG-4MAL. The gel was subjected to alternating periods of small- and large-strain oscillation. Storage (G’) and loss (G’’) moduli are represented by filled squares and open circles respectively. The large-amplitude strain was held constant while the small-amplitude strain was varied between 0.1, shown in gray, and 0.01, shown in blue.

The time required for complete recovery, which is a manifestation of the relaxation time, depends on the size of the stress applied, even in the small amplitude case, as shown in the shaded area of Fig. 5. The relaxation time represents the characteristic timescale for the material to relax from the deformed state to the equilibrium state, and within the KDR framework is rate dependent, as shown in Eq. (2) [55]. Accordingly, the smaller the strain amplitude is for a given frequency, even within the linear regime, the longer the relaxation time will be. We see experimentally that the smaller strain amplitudes require a longer time than the larger to completely recover, supporting the rate-dependence of the relaxation time. The different rates at which the dynamic moduli recover to their pre-deformed state is another indicator of the success of the KDR model’s ability to describe the complex rheology of the granular hydrogels.

## 4. Discussion

The KDR model with brittility accurately describes the complex rheological behavior of the unannealed granular hydrogels we have studied. It is able to account for variation in material chemistry, microgel design, and granular gel formation methodology. The six-parameter model captures both linear and nonlinear properties and accurately predicts steady and transient behaviors. Because of the quality of the description, we can show the rheology of our granular hydrogels displays nonlocal effects whereby rapid elastic strain enhances the acquisition of plastic strain. By describing the rheology with this model, we can also reduce the complexity presented by the different nonlinear rheological protocols to understand granular behavior under applications of arbitrary forces. The success of the model allows us to connect the model parameters to experimental variables. In particular, the model parameters’ dependence on fractional composition (GelMAL vs. PEG-4MAL) highlights the trend of the linear and nonlinear behavior with the increase of PEG-4MAL fraction. The model effectively elucidates the complex interactions between granular populations, providing an opportunity to semi-orthogonally tune fractional composition and material properties.

Few studies have investigated the rheology of mixtures of discrete populations of microgels or similar objects. Di Caprio et al. reported amplitude sweeps for mixtures of norbornene-modified hyaluronic acid microgels with similarly-sized spheroids [56]. They found that mixtures of microgels with spheroids of fractional composition up to 50% microgels presented similar storage moduli to the modulus of the microgels alone. Their system is different to ours, which may be the source of the different observations: they used batch-emulsified microgels with a higher %CV, and the cell spheroids used in their work present little to no overshoot in the loss modulus, indicating the rheological properties of the resultant mixtures may follow different trends than our microgel-microgel mixtures. We did not observe a strong dependence of either the storage or elastic modulus on microgel size, in general agreement with others’ work [57, 58] with the noted exception that Emiroglu et al. did observe a weak dependence of the plateau modulus upon stiff, but not soft microgels. In isolation, microgel polymer fraction was by far the most significant parameter of the individual factors studies. However, the %CV in microgel diameter, and by extension the method of microgel fabrication, was also notable. This agrees with previous results that have investigated the rheology of granular materials, in which monodisperse granular materials possessed greater elastic moduli than polydisperse ones [59]. Batch emulsification is a desirable method for microgel formation due to its high throughput relative to conventional microfluidic devices. These results emphasize the importance of microgel dispersity and suggest that sieving methods to reduce polydispersity are an important tool in leveraging the throughput of batch emulsification while mitigating any need-specific drawbacks of polydispersity.

In the model, flow behavior is characterized by a Herschel-Bulkley viscosity, which includes a yield stress, a consistency index, and an exponent. Additionally, the brittility factor accounts for yielding behavior that ranges from ductile to brittle. In this study, the brittleness of the material was found to be dependent on the polymer selection, evident in the order of magnitude difference in brittility between 4wt% PEG-4MAL and 4wt% GelMAL. By contrast, the brittility was found to be relatively insensitive to microgel diameter, packing density, and microgel size dispersity. The monotonic variation of the brittility factor with fractional composition (PEG-4MAL and GelMAL mixtures) demonstrates a new design parameter; yielding behavior may be tuned by adding or subtracting a “filler” or alternative microgel design in the granular mixture. This may also suggest a way to use brittility as a parameter to trace changes in cell-mediated remodeling within collections of cell-laden hydrogel microgels; we have previously shown that cells embedded in gelatin hydrogels perform significant remodeling that is an essential part of shaping the activity of stem cell cultures [60, 61] and have developed routes to embed these cells in Gel-MAL microgels [23]. Here, rheological methods that reveal brittility may provide a new route to monitor cell-mediated remodeling within granular assemblies.

The yield stress was notably lower in the PEG-4MAL/GelMAL granular mixtures than in monolithic granular hydrogels, which may be due to the mismatch in microgel properties complicating or broadening yielding behavior. Fundamental to the functioning of the model is the idea that rapid elastic strain enhances the acquisition of plastic strain, which implies heterogeneous yielding [43]. The softer particles within a sea of stiffer ones may act as a sacrificial structure that preferentially yields before the stiffer matrix. From our experiments, we also observed that the yield stress was significantly dependent on microgel polymer wt%,packing density and %CV, and little effect on microgel diameter. These results are particularly relevant for applications in which cellular cargo may be co-injected or allowed to invade. This knowledge is important, given that pore size and porosity are known to be critical, instructive parameters in cell-laden granular gels [62].

Since a key motivation for 3D printing is the stability and fidelity of the final product, the linear viscoelastic properties may be a reasonable starting point in material design, with the caveat that immediate post-print crosslinking may mitigate these concerns. Nevertheless, these properties would direct the engineer to select a material and polymer fraction appropriate for the downstream goal. One may also wish to consider yielding behavior if the processing conditions are considered as well as the properties of the printed product. We observed that the manner in which yielding occurred was highly dependent upon the material choice. If more ductile yielding is a key target, one may wish to form the granular gel from a more polydisperse microgel population or add a less brittle microgel population to the granular gel at some fraction. These choices will also affect the rate of self-healing, because of the role the brittility has in the effective shear rate. Lastly, if the shear stresses during flow [30] are a concern – for example, if cellular cargo are present within or between microgels – the material choice and size variability are likely to be important factors. Cell viability might be improved by selecting a polymer (here, PEG-4MAL) that presents lower steady-flow shear stress, by increasing microgel size variability, or potentially by incorporating another microgel population at some fraction in a heterogeneous mixture.

The parameter design space of granular hydrogels is greatly expanded relative to conventional bulk hydrogels. The number of available materials compounded with microgel and granular properties and methods requires any researcher tailoring a complex granular material to their needs to conduct experiments to understand how their material or mixtures will depend upon design parameters. As a result, this work provides a framework which may be used to better understand granular hydrogel rheology, to pursue systematic studies of granular hydrogel design parameters, and to inform design approaches by the broader soft matter and bioengineering communities who wish to print or deliver complex granular materials for a wide range of applications.

## Acknowledgements

Funding sources include the National Institute of Diabetes and Digestive and Kidney Diseases of the National Institutes of Health under Award Number 2 R01 DK099528, the National Institute of Dental and Craniofacial Research of the National Institutes of Health under Award Number R01 DE030491, and the National Cancer Institutes of the National Institutes of Health under Award Number R01 CA256481. The content is solely the responsibility of the authors and does not necessarily represent the official views of the NIH. The authors are also grateful for additional funding provided by the Department of Chemical & Biomolecular Engineering, the Cancer Center at Illinois, and the Illinois Scholars Undergraduate Research Program at the University of Illinois Urbana-Champaign.

## Contributions (CRediT: Contributor Roles Taxonomy[63, 64])

**G.B. Thompson:** Conceptualization, Data curation, Formal Analysis, Visualization, Investigation, Methodology, Writing – original draft, Writing – review & editing. **J. Lee:** Conceptualization, Data curation, Formal Analysis, Visualization, Investigation, Methodology, Writing – original draft, Writing – review & editing. **K.M. Kamani:** Conceptualization, Investigation, Methodology. **N. Flores-Velasco:** Investigation, Methodology. **S.A. Rogers:** Conceptualization, Resources, Funding acquisition, Supervision, Writing – review & editing. **B.A.C. Harley:** Conceptualization, Resources, Project administration, Funding acquisition, Supervision, Writing – review & editing.

