## Supplemental Information for "Granular hydrogels as brittle yield stress fluids"

^1^ Dept. Chemical and Biomolecular Engineering

^2^ Carl R. Woese Institute for Genomic Biology

^3^ Cancer Center at Illinois

University of Illinois Urbana-Champaign

Urbana, IL 61801

* Co-first authors

**Corresponding Author:**

B.A.C. Harley

Dept. of Chemical and Biomolecular Engineering

Cancer Center at Illinois

Carl R. Woese Institute for Genomic Biology

University of Illinois at Urbana-Champaign

110 Roger Adams Laboratory

600 S. Mathews Ave.

Urbana, IL 61801

**Table S1** Descriptive microgel statistics for all screened conditions, and associated method. MFD = flow focusing microfluidic device. BE = batch emulsion. Flowrate refers to the flowrate of the crosslinker emulsion that pinches the hydrogel precursor into droplets in the microfluidic device, while RPM refers to the heating plate RPM used to generate microgels with the batch emulsion process.

| Polymer | Polymer wt% | Method | Flowrate or RPM | Mean (µm) | SD (µm) | CV (%) |
| --- | --- | --- | --- | --- | --- | --- |
| PEG-4MAL | 5% | MFD | 40 µL/min | 201.9 | 7.7 | 3.8% |
|  |  |  | 50 µL/min | 184.1 | 5.9 | 3.2% |
|  |  |  | 60 µL/min | 164.7 | 4.3 | 2.6% |
|  |  |  | 70 µL/min | 153.9 | 13.0 | 8.5% |
|  |  |  | 80 µL/min | 139.7 | 5.7 | 4.1% |
|  | 4% |  | 50 µL/min | 186.6 | 5.6 | 3.0% |
|  |  |  | 60 µL/min | 165.2 | 4.9 | 3.0% |
|  |  |  | 70 µL/min | 156.3 | 15.1 | 9.7% |
|  |  |  | 80 µL/min | 130.0 | 7.7 | 5.9% |
|  | 5% | BE | 200 RPM | 196.9 | 81.1 | 41.2% |
|  |  |  | 300 RPM | 137.2 | 40.8 | 29.7% |
|  |  |  | 400 RPM | 119.4 | 28.8 | 24.1% |
|  |  |  | 500 RPM | 109.2 | 18.7 | 17.1% |
| GelMAL | 5% | MFD | 80 µL/min | 225.9 | 16.4 | 7.3% |
|  | 4% |  | 80 µL/min | 110.3 | 9.7 | 8.8% |

**Table S2** Table shows all values fitted to the KDR model for heterogenous granular hydrogel data depicted in Fig 3, as well as the method/source for parameter determination

|  | Elastic modulus, G (Pa) | Structural viscosity, $\eta_{s}$ (Pa-s) | Yield stress, $\sigma_{y}$ (Pa) | Consistency index, k  (Pa s^n^) | Exponent, n | Brittility factor, Bt |
| --- | --- | --- | --- | --- | --- | --- |
| Gel 100% | 225 | 10 | 135 | 1 | 1.5 | 2 |
| Gel 67%  PEG 33% | 341 | 14 | 90 | 20 | 0.7 | 4 |
| Gel 33%  PEG 67% | 850 | 32 | 79 | -15 | 0.5 | 5 |
| PEG 100% | 1610 | 50 | 160 | -55 | 0.2 | 8 |
| Source | Linear regime frequency sweep | | Nonlinear regime amplitude sweep | | | |

**Table S3** Impact of investigated granular hydrogel parameter changes upon monolithic PEG-4MAL granular hydrogel properties incorporated into the KDR model.

|  | Elastic modulus, G (Pa) | Structural viscosity, $\eta_{s}$ (Pa-s) | Yield stress, $\sigma_{y}$ (Pa) | Consistency index, k  (Pa s^n^) | Exponent, n | Brittility factor, Bt |
| --- | --- | --- | --- | --- | --- | --- |
| Large Diam. | 1690 | 40 | 185 | -9 | 0.95 | 7 |
| High Dens. | 1625 | 40 | 210 | -90 | 0.2 | 8 |
| Batch | 1250 | 42 | 110 | -50 | 0.1 | 5 |
| Low wt% | 925 | 24 | 55 | 0.5 | 0.76 | 20 |
| Source | Linear regime frequency sweep | | Nonlinear regime amplitude sweep | | | |

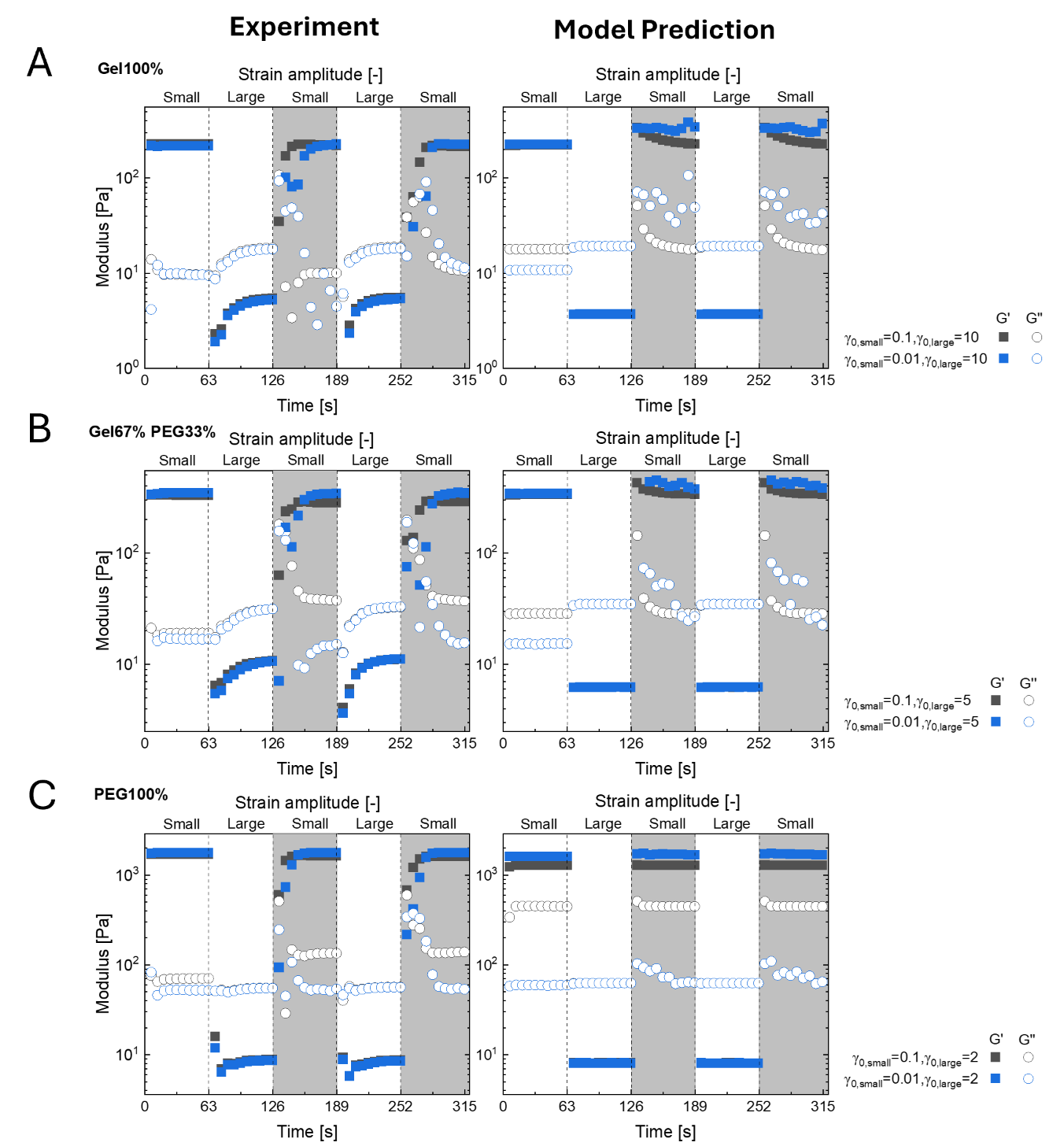

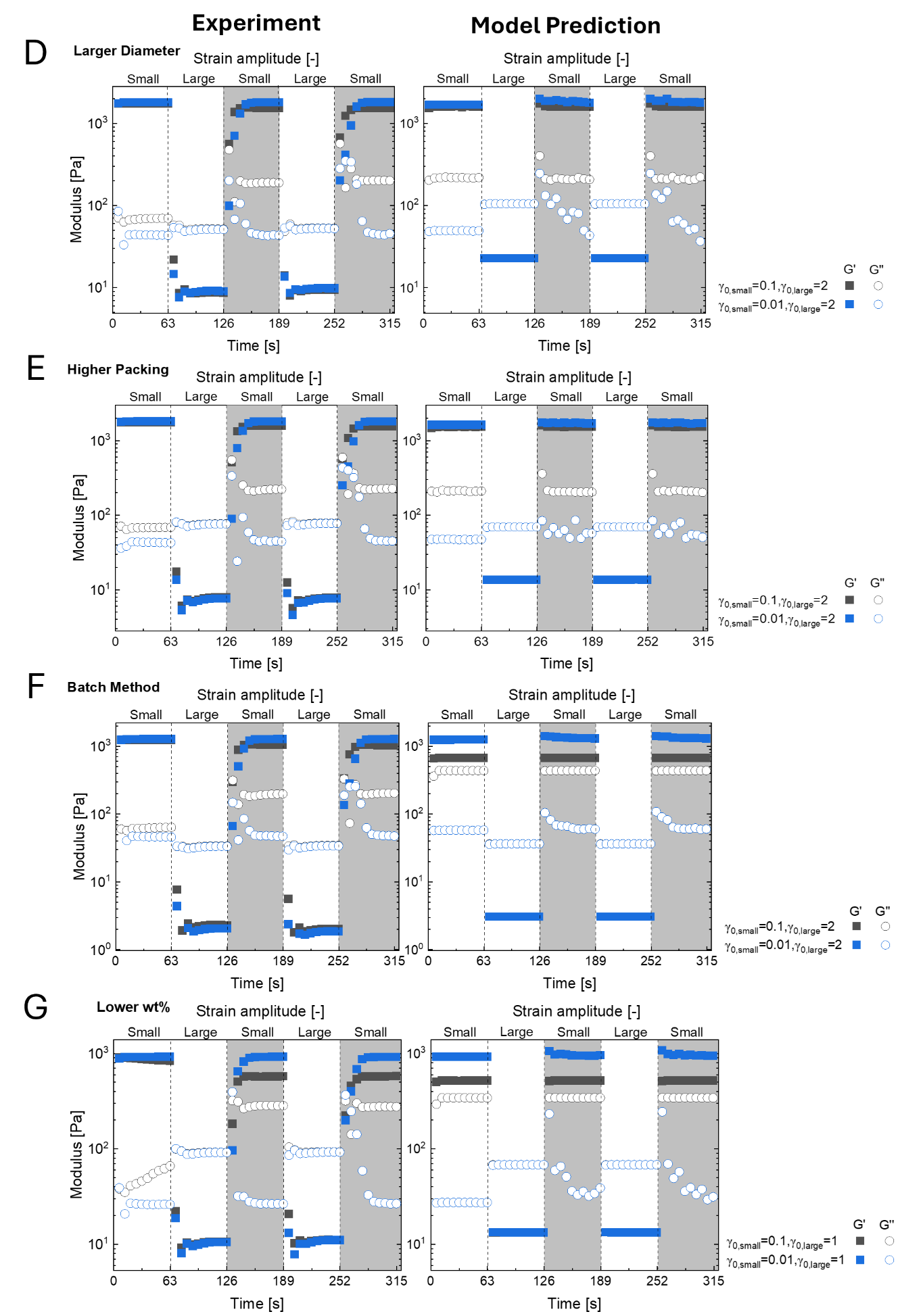

**Figure S1.** Experimental (left) and KDR model-predicted (right) self-healing data for granular hydrogel parameter changes upon **A** 100% GelMAL, **B** 33% PEG-4MAL / 67% GelMAL, **C** 100% PEG-4MAL, **D** PEG-4MAL with larger diameter microgels, **E** PEG-4MAL with higher packing density, **F** PEG-4MA formed from batch-emulsified microgels, and **G** PEG-4MAL with lower-wt% microgels
